# Effects of active and passive enrichment regimes on fecal glucocorticoid metabolite levels in captive Indian leopards (*Panthera pardus fusca*)

**DOI:** 10.1101/2021.12.13.472403

**Authors:** Nirali Panchal, Chena Desai, Ratna Ghosal

## Abstract

Environmental enrichment improves health and wellbeing of zoo animals. To test this hypothesis, we used Indian leopards, one of the popular zoo animals, as a model system to understand effects of active (interacting) and passive (noninteracting) enrichment elements on stress hormone levels of captive individuals. We included three enrichment categories, category ‘A’ (having both active: large size cage, and passive: controlled temperature, playback of forest sounds and sound proof glasses to filter visitors’ noise, enrichment elements), category ‘B’ (active enrichment type I, small size cage with air coolers), and category C (active enrichment type II, medium size cage without air coolers) for the leopards (n=14) housed in two Indian zoos. We standardized a non-invasive method to measure fecal glucocorticoid metabolite (fGCM) levels in captive leopards. The standardized fGCM assay was further validated by analysing samples from free-ranging leopards, as well. The fGCM levels (Mean±SE) were 10.45±2.01 and 0.95±0.003μg/g dry wt of feces in captive and free-ranging leopards, respectively. Our results demonstrated that fGCM levels of leopards in categories B and C were significantly (P<0.05) different from each other, thus, indicating cage size (an active enrichment element) as an important factor in influencing the physiology of the sampled animals. Overall, the findings of the study will contribute towards informing policies for management of the Indian leopards.

## Introduction

Conservation action requires sustainable management of captive and zoo-housed individuals of a species. Globally, there are approximately 10,000 zoos and captive breeding centres, and India alone has more than 145 zoological gardens [1]. Most of the natural populations of animals are declining due to deforestation and urbanization, thus, zoo or captive populations provide an alternative source of genetic materials for vulnerable or threatened species [2]. Further, researchers extensively use zoo animals for behavioural and ecological studies to better understand the wild populations of the target species, and to develop an informed management action plan for the same. Besides conservation research, zoos are also a source of recreation, entertainment and education for the general public [3,4]. Thus, as a part of the ex-situ conservation action plan, monitoring health and wellbeing of zoo animals becomes a topmost priority for ecologists and conservation managers, as well. However, being away from their natural habitat, zoo animals often suffer (mentally and physically) within the artificial, unfamiliar, captive environment [5,6]. Thus, to maintain the wellbeing of the zoo animals, several types of physical and virtual stimuli needs to be added to the captive environment with an attempt to supplement the captivity with enriched habitat conditions.

Enrichment is the alteration made to the environment of captive animals for their physiological and psychological wellbeing. Studies have shown that enrichment can be divided into two categories, active and passive. Active enrichment can be defined as enrichment that requires animals to perform some sort of physical activity or direct interaction with a physical object. For example, a study in literature [7] shows, structural enrichment in the form of proper housing conditions increased the success rate of reproduction in animals like rats, ferrets, great apes and ungulates. Similarly, in case of captive bears, stereotypic behaviour was reduced by food stimulation [6]. Bears had higher locomotor and exploratory behaviors when food was provided in different ways, such as in a log filled with honey and by hiding food throughout the exhibit. Multiple ways of food presentation helped bears to reduce stereotypic pacing from 125min/day to 20min/day, and increased the rate of their exploratory behaviours [6]. On the other hand, passive enrichment can be defined as a regime that does not include any interaction with a physical structure or any kind of direct contact with a living being. Passive enrichment includes visual, olfactory, and auditory enrichment, and mostly consists of modifications that enrich the ambient environment without necessarily involving any physical interaction. Television, video and computer games, mirror and colours are known to be effective as visual, passive enrichment for rhesus macaques, horses and chimpanzees [8]. Whereas in the case of auditory enrichment, sounds specific to animal’s natural environment, and even classical music, radio broadcasts and instrumental music acted as enrichment to female African leopards, Asian elephants, Gorillas, Guinea pigs and rats [8]. A four-month long study on zoo-housed female Asian elephants showed that classical music reduced stereotypic behaviours in elephants when compared to the group that received no auditory enrichment [5]. Management of zoo animals is challenging due to vast differences between the natural and the captive environment, and thus, captive conditions often trigger physiological stress. Long-term physiological stress is highly detrimental to zoo animals leading towards suppression of reproductive and immune systems, and inducing unusual behavioural changes, which may eventually impact an animal’s wellbeing.

Stress can be defined, as a condition of loss of homeostasis in the body and stressor is an event or force, which causes this disruption. A series of physiological events that take place to restore this disruption is defined as a stress response. Stress can be physiological and psychological, as well. Physical stressors, for example injuries, conflict [9], environmental factors such as temperature, humidity, sunlight, as well as internal factors such as anoxia, hypoglycaemia, contribute to physiological stress [10]. Similarly, stimuli affecting the emotions such as anger, excitation, fear, anxiety, can cause psychological stress [10]. Under stressful conditions, stress hormones are released as a biological response helping to cope with “fight or flee” situations. Most kinds of stressors trigger the release of glucocorticoid (GC) hormones [10], and GCs are known to increase blood glucose levels, suppress immune system to help maintain homeostasis, and support metabolism of fats, proteins and carbohydrates [11,12] to fight against the stressful situations. Studies have shown that captive conditions are highly stressful for most animals and often, such artificial environment may lead towards poor reproductive performances and/or higher prevalence of diseases among the zoo populations [13,14].

Confinement, artificial environment, visitors [15] and isolation of social animals [16] are the key factors known to cause high levels of stress in captive animals. Thus, to improvise ex-situ conservation efforts, enriched habitats are provided to captive animals and their physiological response towards the enrichment can be assessed through measurements of stress hormones, mostly by monitoring the levels of GCs [17,18]. For example, a study on small felids showed that female cats when provided with bigger enclosures, an example of active enrichment, had reduced stress hormone levels, and resumed reproductive cyclicity when compared to females maintained under non-enriched, stressful conditions (small and barren enclosures) [17]. Another study on Clouded leopards showed that certain unusual behaviours, for example, fur plucking and nail biting, were relatively higher in captive animals, thus, indicating stressful conditions [19]. To reduce such behaviours, different types of enrichment (active and passive) were provided to captive Clouded leopards [19], of which increased time spent with the keeper, a type of active enrichment element, was found to be effective in reducing the high faecal glucocorticoid metabolite (fGCM) levels in Clouded leopards. Globally, a large population of different species are present in captivity, and with an aim to provide improvised husbandry practices; studies need to be conducted on understanding the effect of diverse enrichment elements (active or passive or a combination of both) on the physiological wellbeing of captive populations.

India is home to rich biodiversity and hosts a large population of captive animals, as well. Monitoring wellbeing of the captive animals is one of the priorities for the managers and the veterinarians of the Indian zoos. One such popular animal housed in a large number of Indian zoos is the Indian Leopard, categorized as endangered as per the IUCN Red List (2016). Out of 145 zoos in India, leopards are present in 76 different zoos across the country [20], and each zoo having approximately 8-10 leopards. Studies have shown that captivity is very stressful for leopards and induces all sorts of unusual behaviours. For example, leopards kept in zoos in 3 different states of India (Maharashtra, Kerala and Delhi) showed high levels of stereotypic behaviours (repetitive walks, chewing paws and snapping) and had higher faecal glucocorticoid (fGC) levels under conditions with no enrichment [21]. However, fGC levels and intensity of stereotypic behaviours reduced when animals were provided with different types of structural enrichment, for example, den or pool within an enclosure, and more time spent with a keeper providing care. Another study showed that captive leopards preferred to spend more time in enriched zones having structural elements, for example, trees, barrels, sleeping platform and logs, and had decreased their rate of stereotypic pacing under such conditions [22]. However, both the studies on the captive Indian leopards included components of active enrichment only. To the best of our knowledge, no study has been conducted, so far, to assess the effects of passive enrichment components (auditory, visual or olfactory enrichment) on the physiological wellbeing of the captive Indian leopards, and thus, warrants future investigation.

In this study, we monitored the impact of three different enrichment conditions (referred as categories ‘A’, ‘B’ and ‘C’) on the stress physiology of the zoo leopards. Category ‘A’ had both active and passive elements whereas category ‘B’ and ‘C’ (active enrichment type-I and type-II, respectively) had only active enrichment elements. This is the first study to include both active and passive enrichment elements and assess their effects on the physiological stress levels of the Indian leopards. To measure physiological response towards a particular enrichment regime, we standardized an assay method to measure fGCM levels in captive Indian leopards. For assay validation purposes only, fGCM concentrations were also measured in scat samples collected from free-ranging leopards, as well. The overarching goal of our study is to develop an improvised protocol for better management practices for the target species.

## Materials and Methods

### Study site and animals for captive sampling

Sampling was carried out for 14 leopards (8 males and 6 females) belonging to two zoos of Ahmedabad and Baroda cities within the state of Gujarat, India (Figure 1). Four male leopards and four females were from Kankaria zoo, Ahmedabad, located in the southeastern part of the city, while remaining four males and two females were from Sayajibaug zoo, Baroda. The sampled leopards were either captive-born or captured from the wild, and time since captivity varied from 3 to 18 years. Table 1 presents detailed information (details on rescue, current age, and breeding history) on each of the sampled leopards in both Kankaria and Sayajibaug zoos. All necessary permits and ethics approval were obtained from the concerned authorities.

**Table 1:**
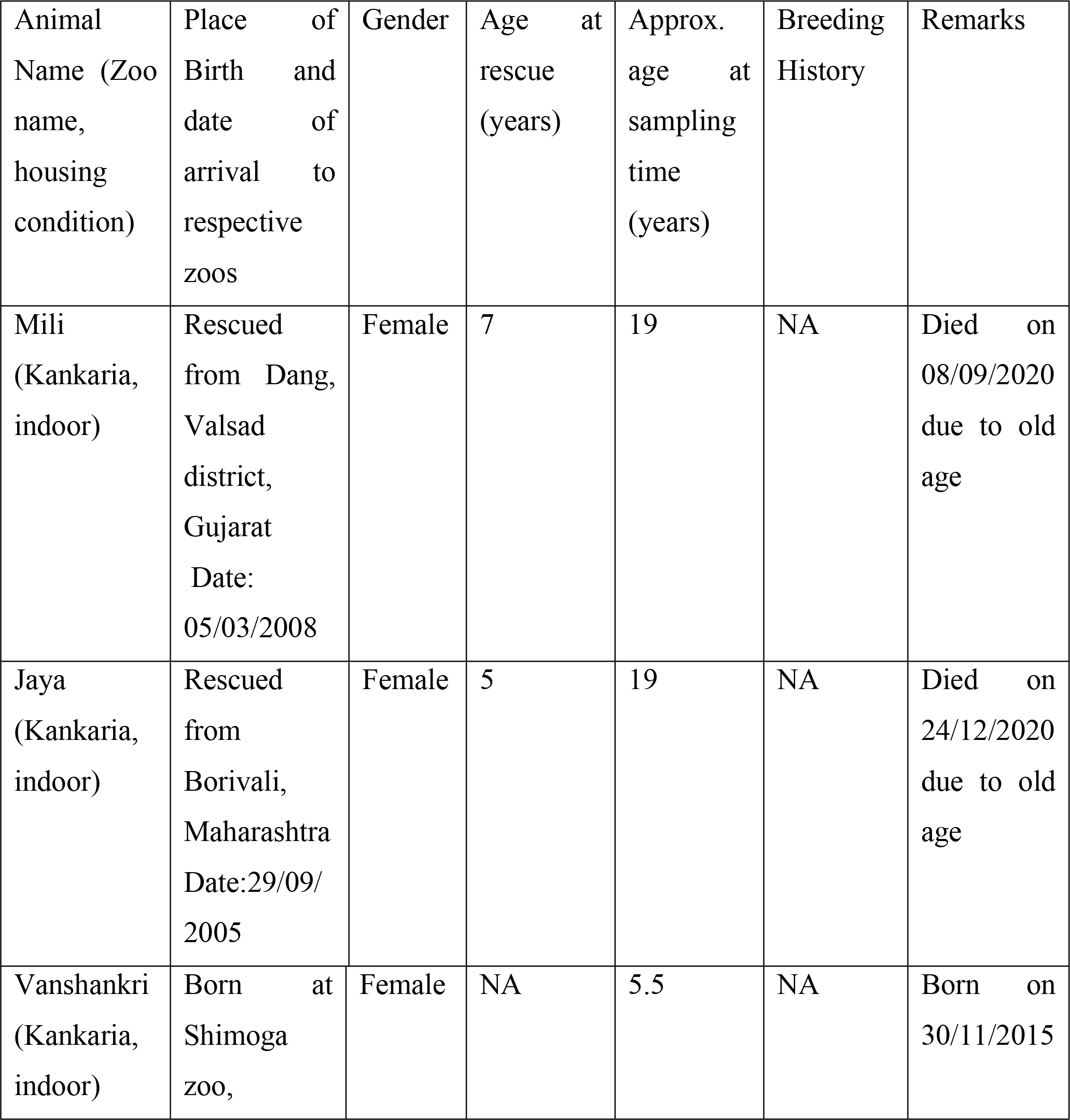
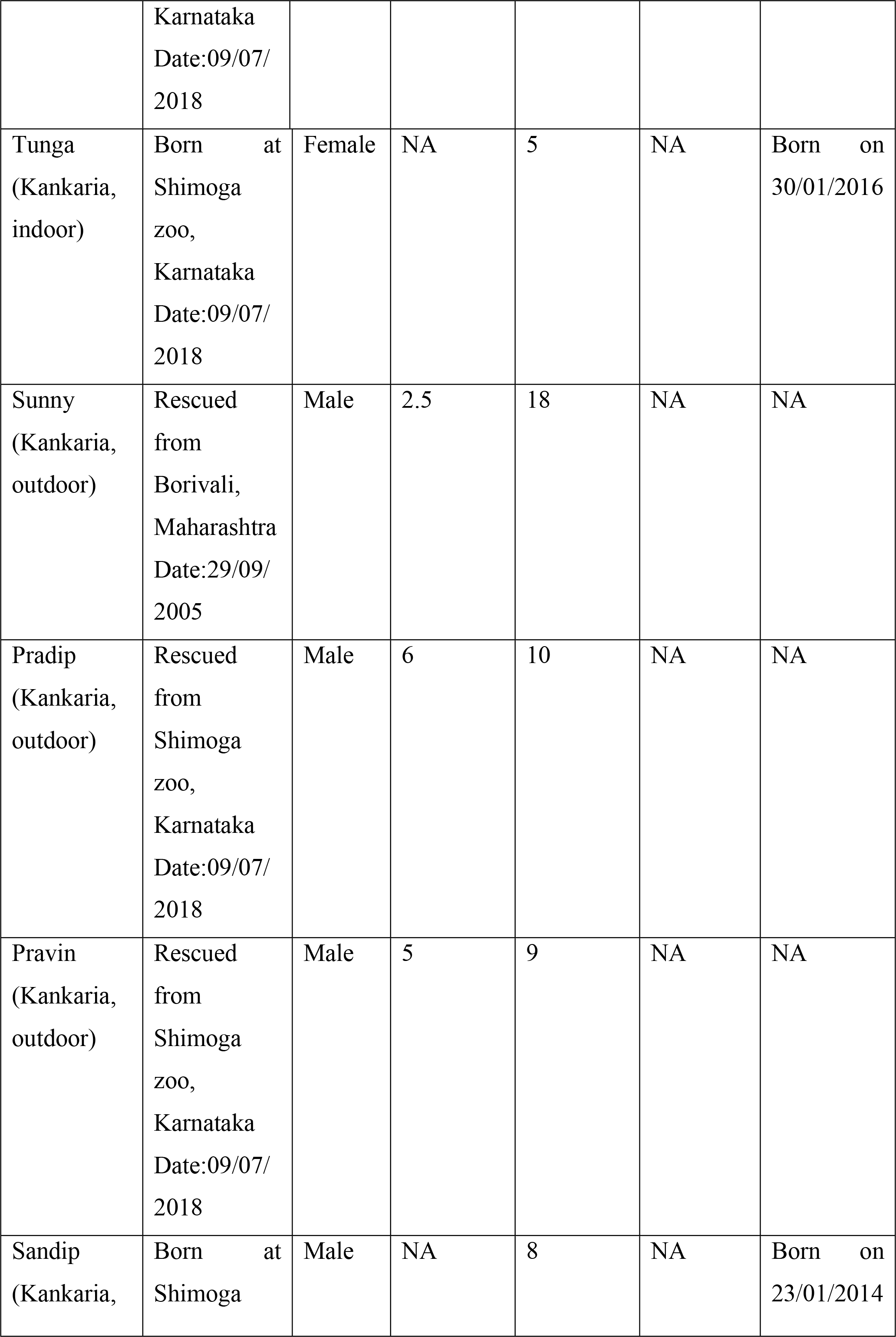
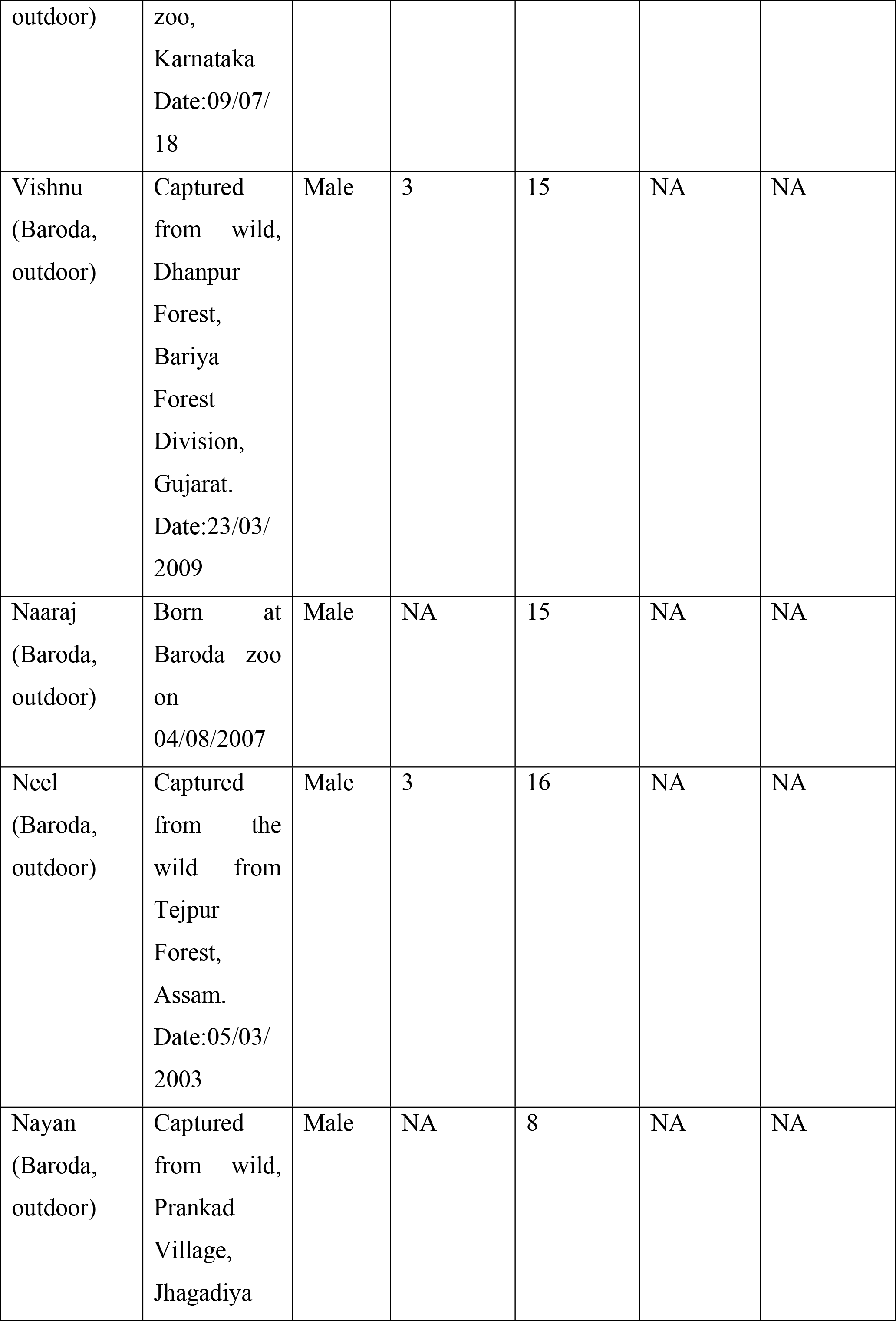
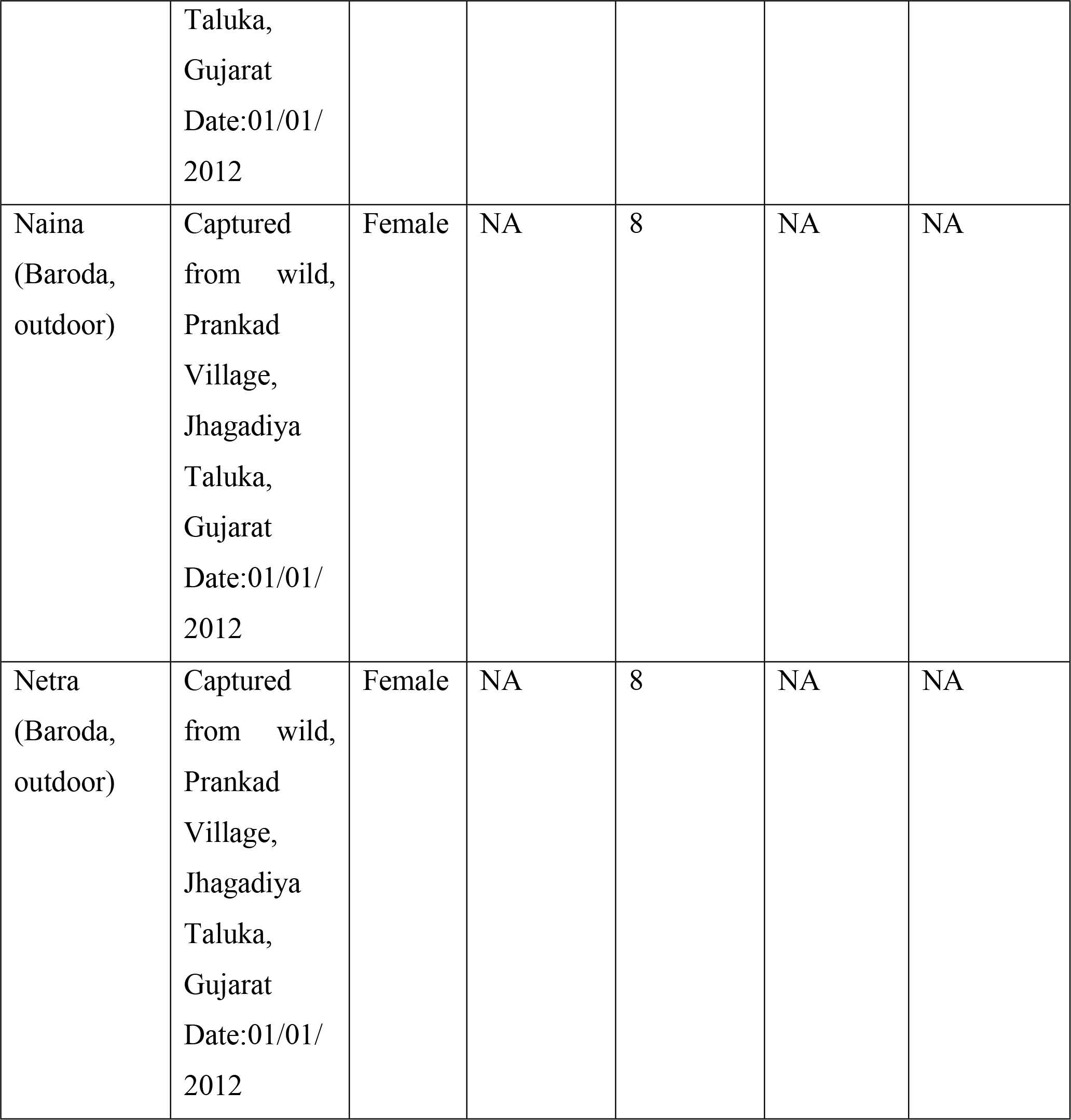
Detailed information on sampled individuals at both Kankaria and Baroda zoos, Gujarat

**Figure 1:**
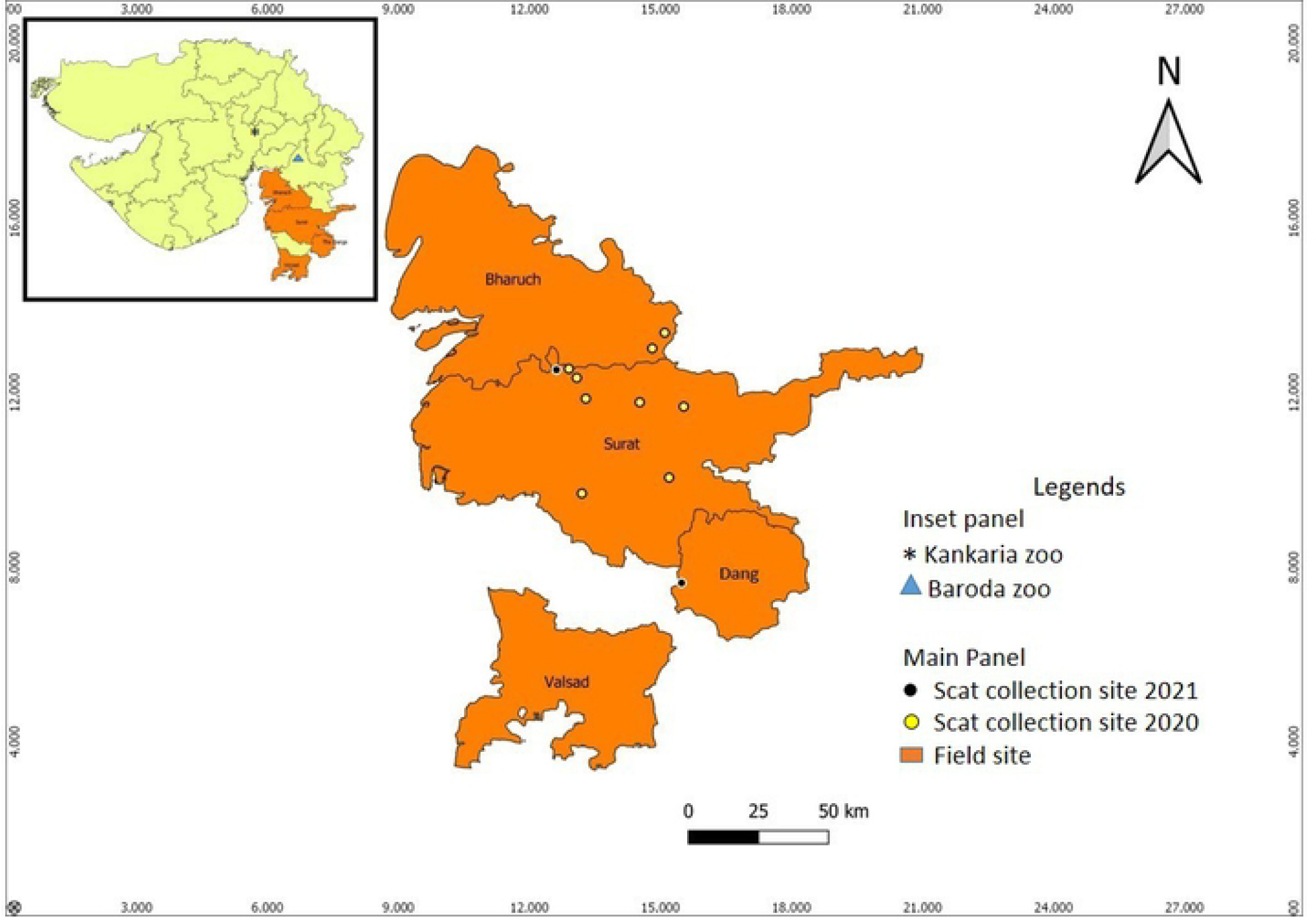
Map showing the study sites for captive and free-ranging populations of the Indian leopards. Inset panel shows the entire map for the state of Gujarat, India, with approximate locations of Kankaria and Baroda zoos, the study sites for captive leopard populations. Main panel shows the scat collection sites for free-ranging leopards in South Gujarat.

### Description of enrichment regimes for captive leopards

Kankaria/Kamala Nehru zoo in Ahmedabad city, Gujarat, is spread over an area of 117 hectares housing hundreds of species of mammals, reptiles and aves. It had two different housing conditions for mammals (indoor and outdoor, Table 2). Sayajibaug zoo (referred to as Baroda zoo from here onwards) in Baroda city, Gujarat, inhabits a total of 874 animals belonging to 89 different species. Baroda zoo had only one type of housing condition, the outdoor (Table 2). Both the zoos are only 72 miles apart from each other, and experience a similar weather pattern in terms of temperature and rainfall. Leopards in both the zoos were adults (Table 2) and were maintained under a similar diet regime. From now onwards, we will refer to the housing conditions in two zoos as category ‘A’ for Kankaria indoor, category ‘B’ for Kankaria outdoor, and category ‘C’ as Baroda outdoor. Table 2 gives details on active and passive enrichment elements that were provided to the leopards within each of these categories, and the number of leopards maintained under these conditions. In all the three categories, leopards were housed in individual cages. Since leopards use height of a cage for climbing and jumping activities, we calculated cage size as a product of length, breadth and height of the cage. Category ‘A’ had the highest enrichment, having the largest (1204 m^3^) cage size when compared to other two categories, and had several active (earthen flooring, larger cage, raised platforms) and passive (sound proof glass to filter visitors’ noise, controlled temperature and playback of natural, forest sounds) elements for enrichment (Table 2). Category ‘A’ was the only regime that had passive enrichment elements for the leopards. Out of all the three categories, category ‘B’ had the smallest cage size (264 m^3^) and was provided with a few active, structural enrichment elements, for example, air coolers during summer, earthen flooring and raised platforms for climbing. Category ‘C’ had medium size cages (517 m^3^) but had no coolers, and was provided with similar enrichment elements as category B, having earthen floors and raised platforms (Table 2).

**Table 2:**
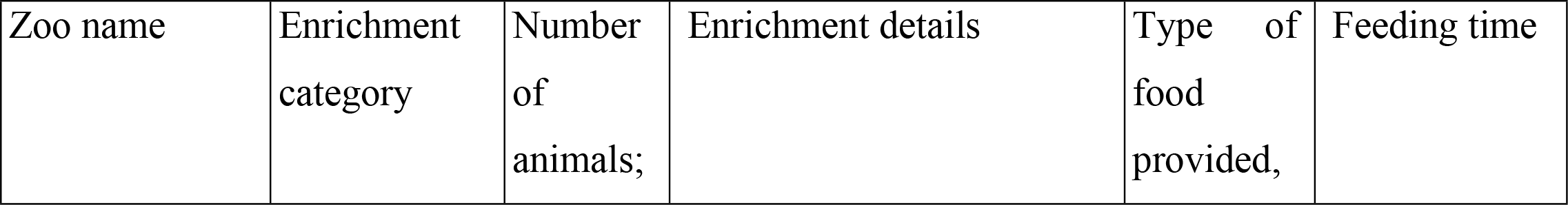
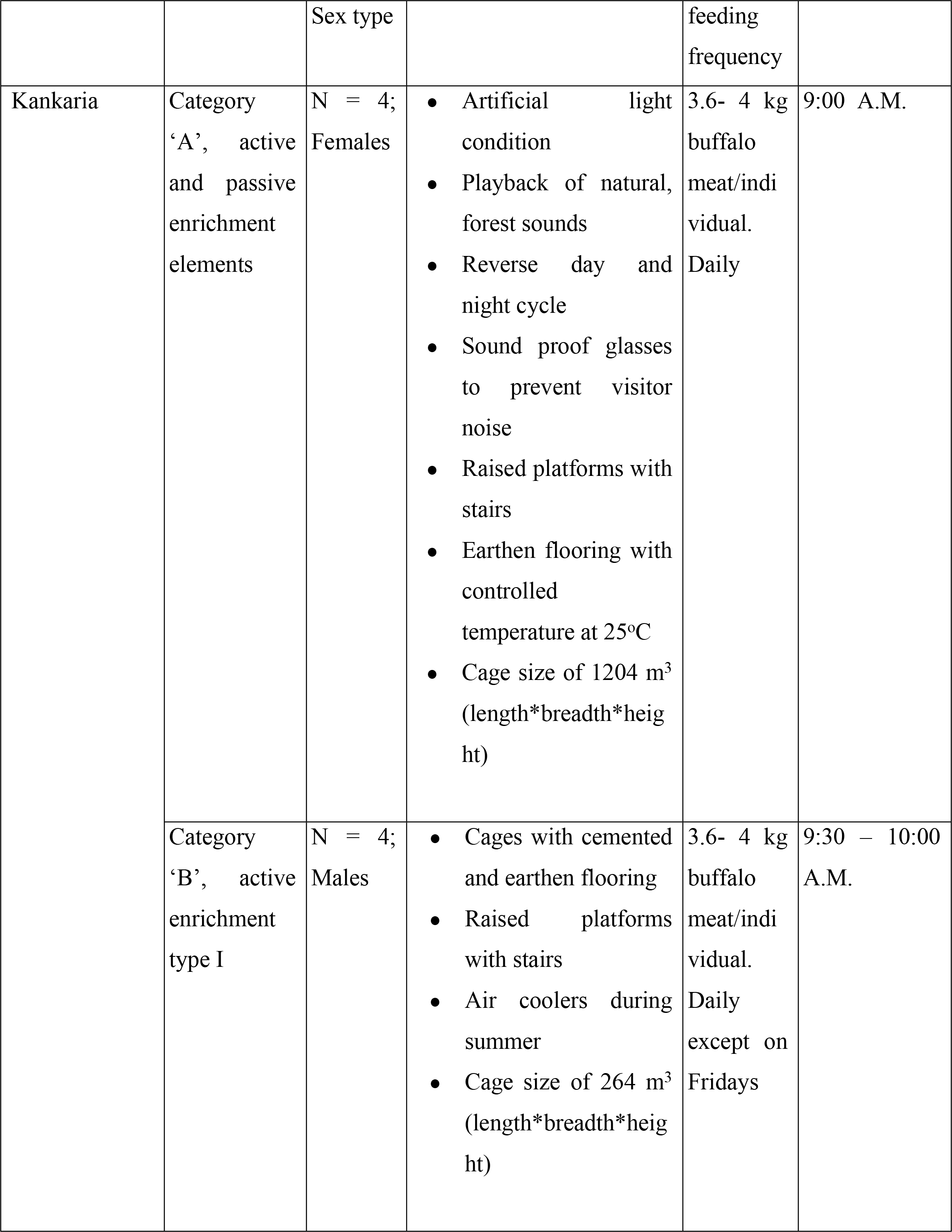
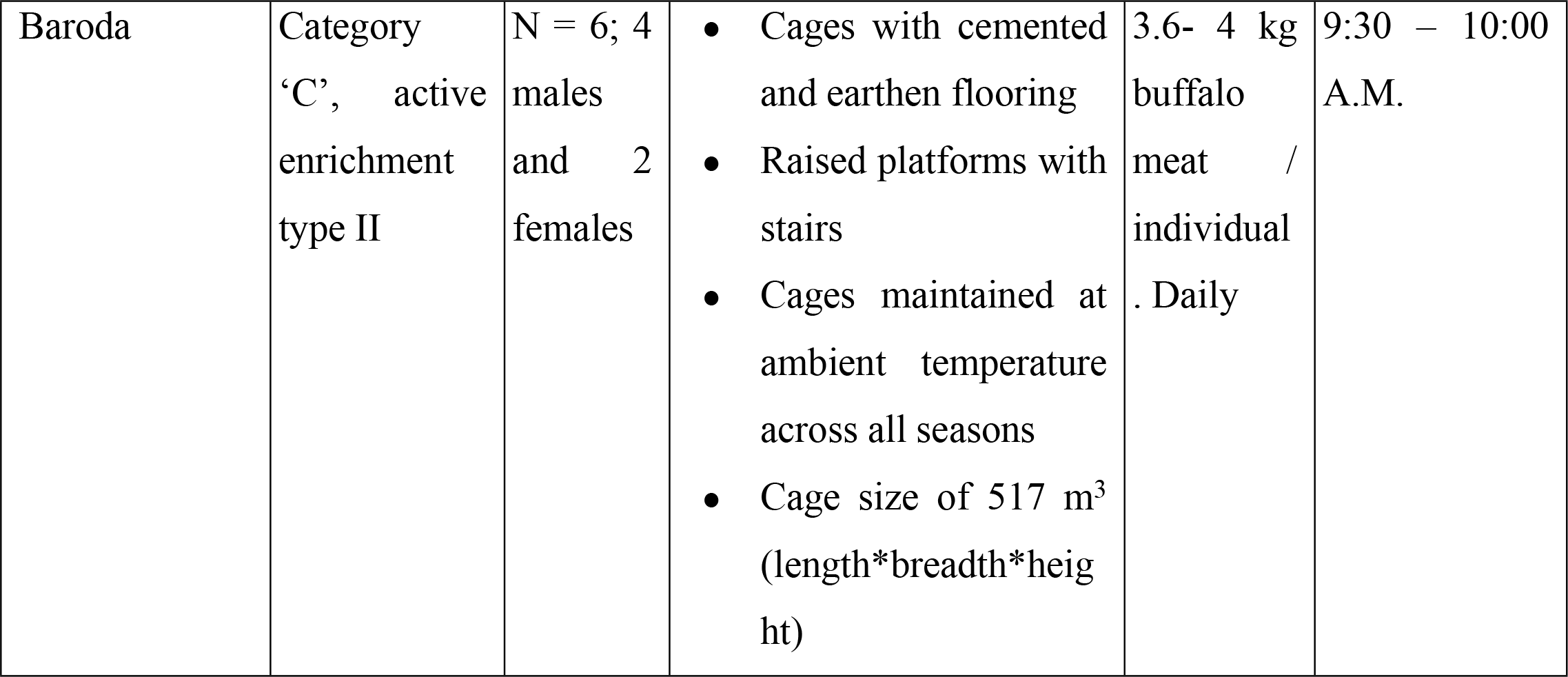
Details on enrichment regimes for leopards in Kankaria and Baroda zoos, Gujarat

### Fecal sample collection from zoos

For both the Kankaria and Baroda zoos, summer sampling (with ambient temperature ranging from 39-42ºC) [23] was conducted from June to July 2019. Similarly, for both the zoos, winter sampling (with ambient temperature ranging from 28-31ºC) [24] was carried out from December to February 2020. Fecal samples were collected twice a week from all the leopards across the three categories, namely, ‘A’, ‘B’ and ‘C’. In total, 119 samples were collected from 14 leopards (n=68 samples for males and n=52 samples for females) across the three categories. Supplementary Table 1 provides description on sampling details with respect to each of the categories (‘A’, ‘B’ and ‘C’) for both the summer and the winter seasons. Due to defecation in water or defecating at an inaccessible spot within the cage, we were not able to collect samples from a few individuals during certain weeks. Samples (∼5 g) were collected using ice-cream sticks and zip lock bags. After collection, samples were stored at −20**°**C until analysis.

### Study site for sampling free-ranging leopards

Four districts, namely, Surat, Bharuch, Valsad and Dang within southern part of Gujarat state, India, were chosen as the field sites for sampling scats from free-ranging leopards. The annual rainfall in south Gujarat is 900-2800 mm, and the average maximum temperature in summer goes up to 40ºC in May which is the hottest month whereas minimum temperature lies at 26ºC. December is the coldest month of the year, and the maximum temperature averages to 25ºC and the minimum is 16ºC [25]. South Gujarat has the highest forest cover with dense canopy. Dang district has a forested area of about 1,368 sq. km, Surat has an area of 496.72 sq.km, Bharuch with an area of 1142 sq.km and Valsad having a forest cover of 274.69 sq.km. [25] According to the census in 2016, estimated leopard population in Gujarat was about 1395 [26]. Out of which, the population count was 43 in Surat, 18 in Valsad and 43 in Dang districts of Gujarat, however, the population count for Bharuch was not available. This accounts for 7.4% of the total leopard population of the Gujarat state [26].

### Fecal sample collection from wild

Scat samples were collected from Surat, Bharuch and Dang districts of Gujarat during January-March 2020, and during the month of March 2021. No scats were obtained from Valsad district. A total of 10 field hours were spent daily in tracking and collecting samples of the leopards. Only fresh scat samples were collected in a zip lock bag. Parameters like moisture content, smell and state of decomposition were used as a criteria to determine freshness of the scat [27]. Scats were kept in an ice-box at the field site and were transferred to −20ºC within 4-6 hours after collection. A total of 12 samples were collected randomly from fairly distant areas (80 to 140 km) across three districts of Gujarat. This was done to avoid pseudo replication by collecting repeated samples from the same individual. However, due to random sampling, the life history or the gender of the sampled leopards could not be determined. Figure 1 shows the sampling sites for free-ranging leopards within the state of Gujarat.

### Extraction of steroid metabolites from fecal samples

Samples were dried using a hot air oven for 24-36hr, until completely dry. After drying, the samples were pulverized and sieved, and the powdered sample was stored in glass bottles at room temperature until further use. For sample extraction, 0.1g of fecal powder was added to 3ml of 80% methanol. The mixture was vortexed for 3 minutes and then centrifuged at 1500 rpm for 10 minutes [28]. Post centrifuge, supernatant was collected (1.5-2ml) in tubes and stored at −20ºC until hormone analysis.

### fGCM hormone assay

fGCM was quantified using ELISA kit purchased from Dr. Rupert Palme in the Department of Biomedical Sciences/Physiology, University of Veterinary Medicine, Vienna. [9] The kit was already validated (ACTH challenge was performed) for African leopards (*Panthera pardus pardus*,) [9]. The kit measures 5α-pregnane-3β,11β, 21-triol-20-one, which is one of the predominant glucocorticoid metabolites in the fecal samples of the leopards. To reduce non-specific binding, the assay was performed on anti-rabbit-IgG-coated (R2004, Sigma-Aldrich) microtiter plates. The antiserum in the kit was against 5α-pregnane-3β,11β,21-triol-20-one (linked at position C20 to carboxymethyloxim) and coupled with bovine serum albumin. The same steroid was used as label, linked at C20 to biotinyl-3,6,9-trioxaundecanediamin [9]. The assay sensitivity was 0.8 pg. The inter and intra assay coefficient of variation (CV) were 14.20% and 9.71% respectively for an internal control sample, whose value lied very close to the EC_50_ of the standard curve. For parallelism, we selected one sample from captivity and one from wild, whose fGCM concentrations were very close to the EC_50_ value of the standard curve, and the fecal extracts were serially diluted and assayed using the same protocol.

### Statistical analyses

To assess the effects of enrichment and season on fGCM values of captive leopards, we used linear mixed effects (LME) model [29] with maximum likelihood method. For the fixed effects, season (with two levels, summer and winter) and enrichment (with three levels, categories ‘A’, ‘B’ and ‘C’), and sex (with two levels, male and female) were included as independent variables, and fGCM concentration was included as a dependent variable. Log-transformed fGCM values were included in the LME model to meet normality assumption. Each individual leopard was repeatedly sampled over time for both the seasons (summer and winter), and thus, individual identity was included as a random effect within the mixed effects model. Model comparisons were conducted to arrive at the best-fit model for the given data set and post-hoc comparisons were done to explore significant interactions within the model. Out of 14 leopards in both the zoos, only four were born in captivity under zoo conditions and the other 10 were captured from the wild (Table 1). Due to disproportionate sample size between captive and wild born leopards, we did not include history of individuals as an independent variable in the model. Time since captivity was also not included in the model as all the individuals spent more than 5 years in captivity, and there were no recent (<1 year) captures. Further, all the sampled individuals were adults (Table 1), thus, age was also not included as an independent variable for the mixed effects model. Adult age criterion for the leopards was followed as outlined by [30].

All analyses were conducted on R version 3.2.3 using ‘nlme’, ‘multicomp’ and ‘ggplot 2’ packages [31]. Post hoc multiple comparisons using Tukey’s HSD were done using the R package ‘emmeans’. Parallel displacement between the standard curve and serial dilutions of the fecal extract was used to validate the fGCM assay for the Indian leopards. The values within the linear range of the curve were subjected to linear regression analysis (PRISM software, version 9) using log molar concentration vs. percent antibody binding of the standard and the sample dilution curves separately. The slopes of the regression lines were compared using Student’s t-test. All the data are reported as Mean± SE and significance level was kept at P<0.05 for all the analyses.

## Results

### fGCM profiles of captive leopards

Overall, the fGCM levels for captive leopards were 10.45±2.01 μg/g dry wt of feces, with male and female leopards having an average value of 10.68±2.96 and 10.13±2.71 μg/g dry wt of feces, respectively. During winter season, the fGCM level, pooled across all the categories, was 18.03±14.68 μg/g dry wt of feces, with individuals maintained in category ‘A’ having a value of 25.11±29.39 μg/g dry wt of feces, those in category ‘B’ showed a value of 27.83±33.26 μg/g dry wt of feces and in category ‘C’, the fGCM level was 1.15±0.59 μg/g dry wt of feces (Figure 2). Meanwhile, during summer season, the overall fGCM level was 8.80 ± 11.98 μg/g dry wt of feces with leopards in ‘A’, ‘B’ and ‘C’ categories had values of 2.42 ± 2.09, 22.62 ± 31.61 and 1.36 ± 0.70 μg/g dry wt of feces, respectively (Figure 2). Figure 3 represents individual fGCM profiles for each leopard during summer and winter months across three categories, category ‘A’ (Fig 3A), category ‘B’ (Fig 3B), and category ‘C’ (Fig 3C).

**Figure 2:**
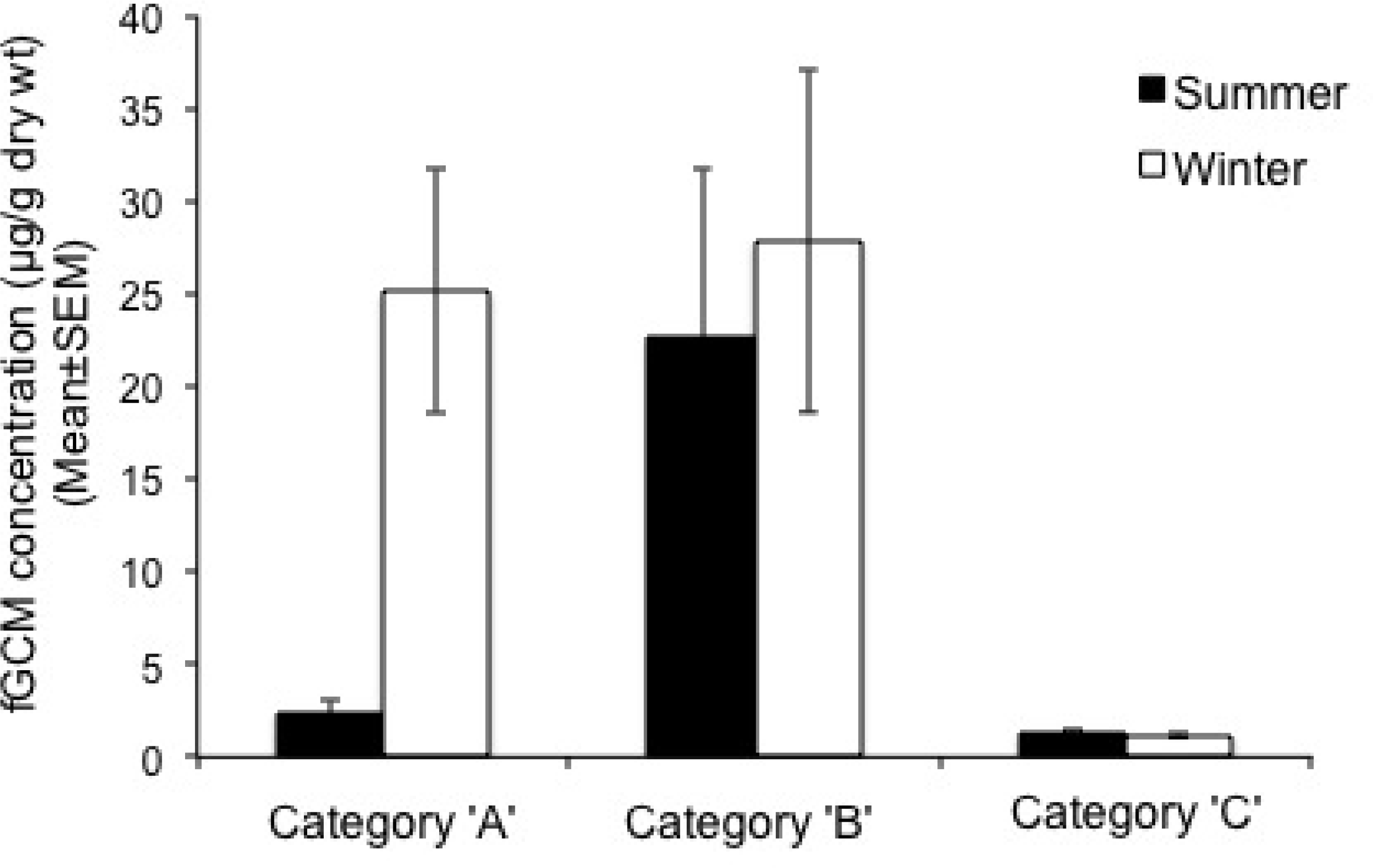
Measured fGCM levels (Mean±SEM) in captive Indian leopards maintained under three different enrichment categories, ‘A’, ‘B’ and C, during winter and summer seasons.

**Figure 3:**
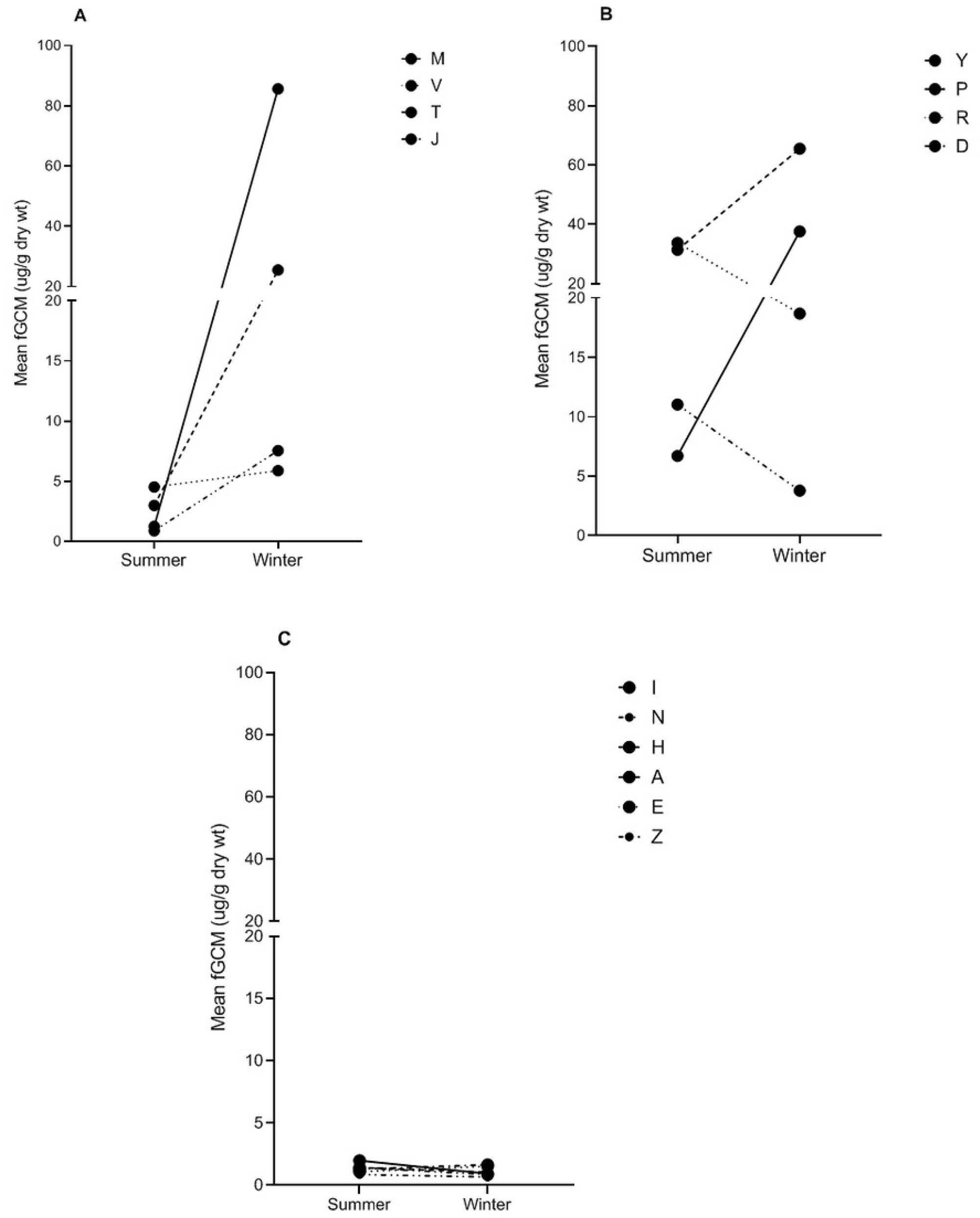
Measured fGCM levels (Mean) in individual captive Indian leopards maintained under three different enrichment categories, ‘A’, ‘B’ and C, during winter and summer seasons. Panel A represents enrichment category ‘A’, panel B represents enrichment category ‘B’, and panel C represents enrichment category ‘C’. A letter of the alphabet represents an individual.

### Effects of enrichment and season on fGCM in captive leopards

A parent LME model was constructed using logfGCM∼Enrichment*Season*Sex type. However, sex type was not significant (P>0.05, data not shown) and was subsequently excluded from the model. Sex type did not produce significant results even after including in two, separate, two way-interaction models, enrichment*sex and season*sex. Further comparison of LME models, with (logfGCM∼Enrichment*Season) and without an interaction (logfGCM∼Enrichment+Season) between enrichment and season, showed significant difference (P<0.001) (Table 3), and thus, interaction term was retained in the final LME model (Table 4).

**Table 3:** Comparison of Linear Mixed Effects model with and without an interaction term

|  | df | AIC | BIC | logLik | Test | L.Ratio | p-value |
| --- | --- | --- | --- | --- | --- | --- | --- |
| Model1<sup>a</sup> | 8 | 341.07 | 363.30 | -162.5358 |  |  |  |
| Model2<sup>b</sup> | 6 | 366.79 | 383.46 | -177.3966 | 1 vs 2 | 29.72157 | <.0001 |

<sup>a</sup>Model1: lme(logfGCM~Enrichment\*Season, random=~1|Ind.ID);
<sup>b</sup>Model2: lme(logfGCM~Enrichment+Season, random=~1|Ind.ID)

**Table 4:**
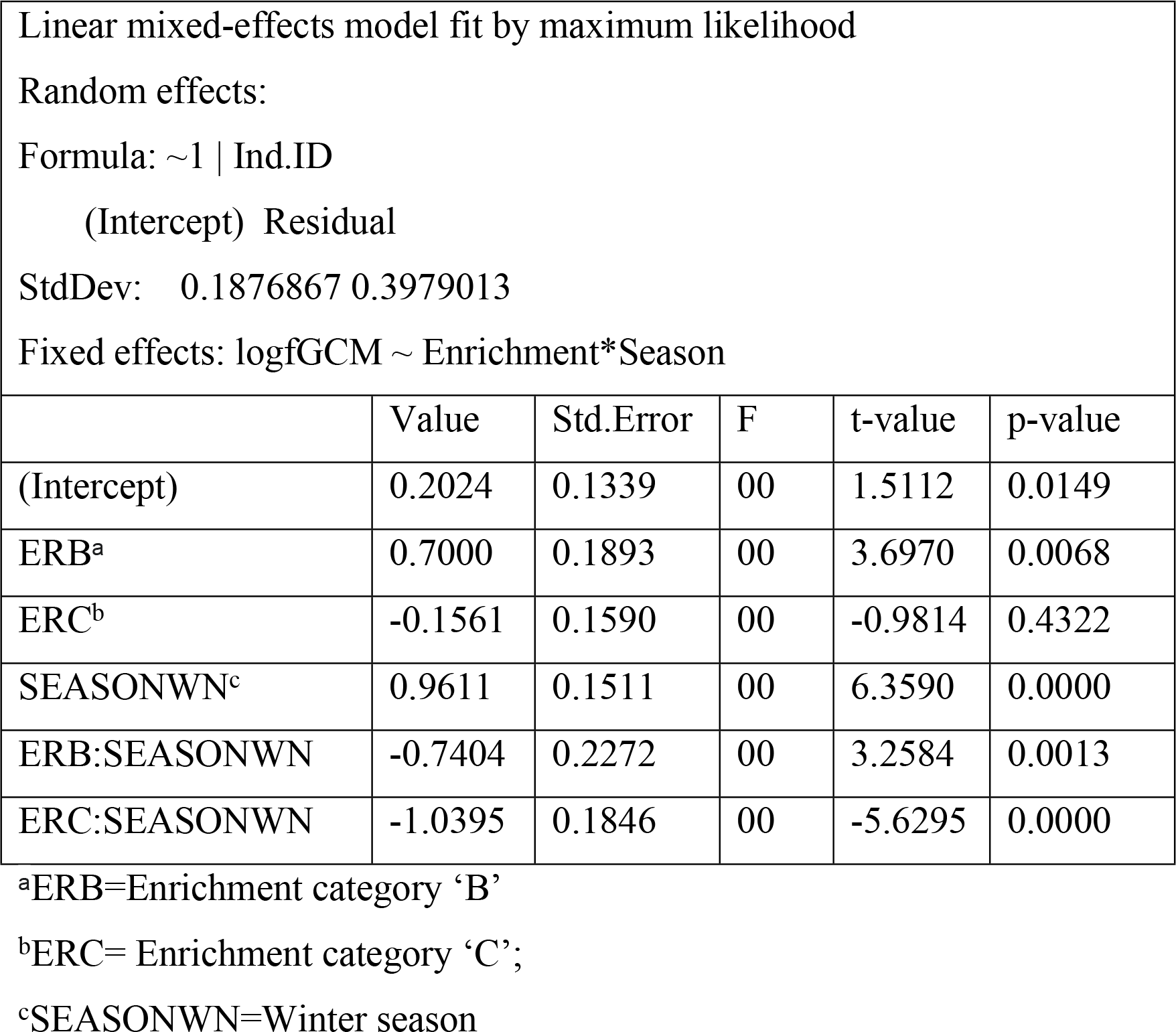
Results from linear mixed effects model showing effects of enrichment categories, seasons and their interaction on fGCM levels in captive leopards

Post-hoc Tukey’s test across different enrichment categories showed significant differences in overall fGCM (pooled across seasons) concentrations between categories ‘A’ and ‘C’ (P<0.001) and between ‘B’ and ‘C’ (P<0.001), as well. No significant differences (P>0.05) were obtained between categories ‘A’ and ‘B’. Further, there was a significant difference in overall fGCM levels (pooled across enrichment categories, ‘A’, ‘B’ and ‘C’) between summer and winter (P<0.01). Within group comparisons (grouped by enrichment category) of pairwise interactions showed significant differences in fGCM levels (Tukey post-hoc test, P<0.01) between summer and winter seasons for leopards in category ‘A’, and no significant differences (P>0.05) in fGCM levels between summer and winter seasons for both categories ‘B’ and ‘C’. When grouped by season, Tukey’s post-hoc test showed significant differences in fGCM levels between categories ‘A’ and ‘B’ (P<0.001), and categories ‘B’ and ‘C’ (P<0.001) during summer season, but no significant difference (P>0.05) was obtained between category ‘A’ and ‘C’ during the summer. In contrast, during winter season, significant differences (Tukey post-hoc test, P<0.001) in fGCM levels existed between categories ‘A’ and ‘C’. Significant difference (Tukey post-hoc test, P<0.001) was also obtained between ‘B’ and ‘C’ category-leopards during the winter season; however, there was no significant difference (Tukey post-hoc test, P>0.05) between ‘A’ and ‘B’ category-leopards for the winter season.

### fGCM profile of free-ranging leopards

A total of 12 samples were collected from three districts of Gujarat. Overall, the fGCM level for free-ranging leopards was 0.95±0.003 ug/g dry wt of feces. The highest and the lowest fGCM values were 0.20 and 1.93 ug/g dry wt of feces, respectively.

### Parallelism curves for captive and wild samples

The slope of the regression lines for standard curve and the serially diluted samples (both captive and wild) (Figure 4) were not significantly different (P>0.05) and thus, the assay can be used to measure fGCM concentrations in both captive and wild Indian leopards.

**Figure 4:**
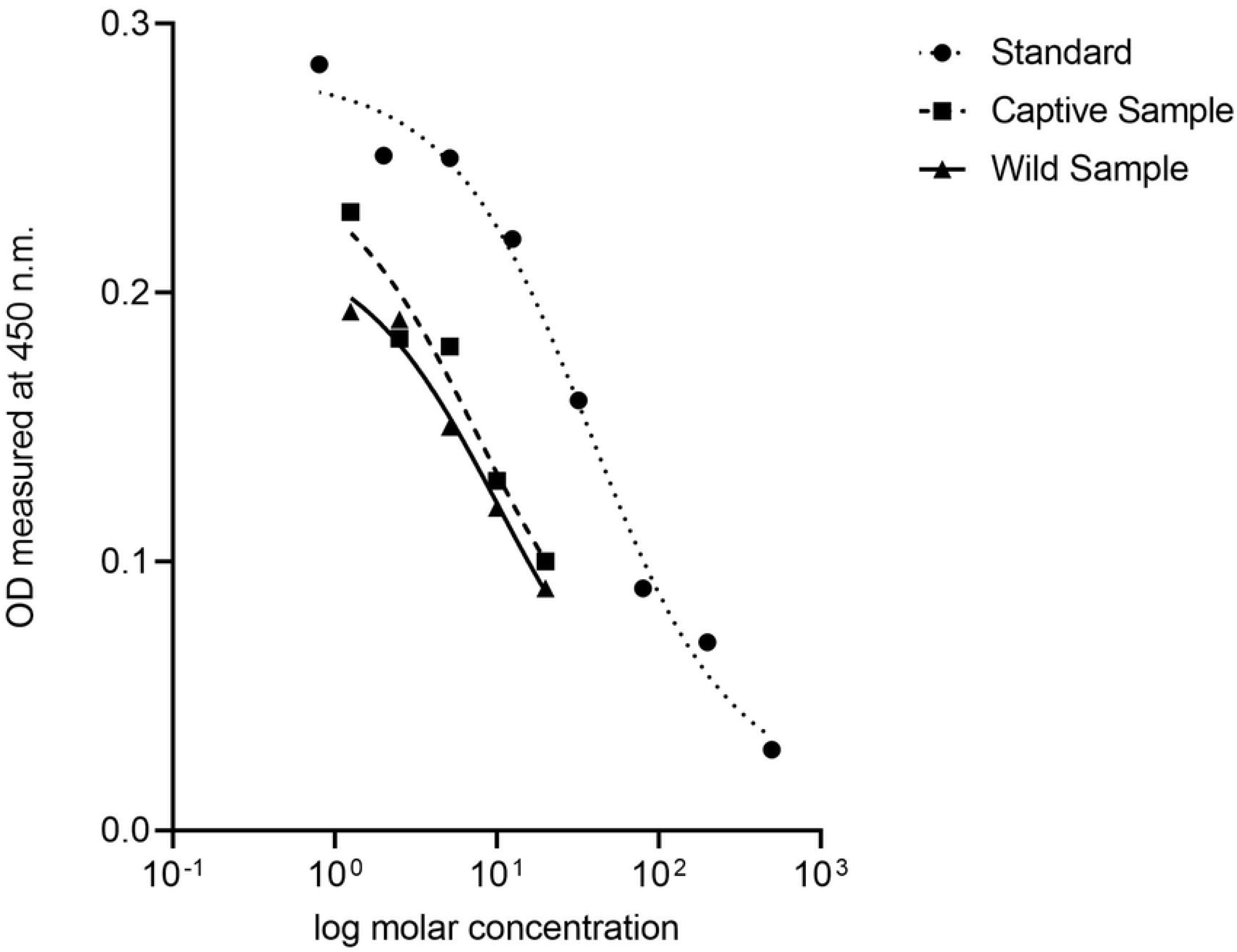
Parallelism of serial dilution of the fecal extract, both wild and captive, with the standard curve.

## Discussion

Our study presents fGCM levels in Indian leopards housed in two Indian zoos and maintained under three different types of enrichment regimes, defined as categories ‘A’, ‘B’ and ‘C’. Overall, the fGCM levels measured in the current study were higher when compared to the levels measured by Vaz et al. (2017) [21], which measured native hormone (corticosterone) levels in the scats of captive Indian leopards rather than metabolite levels. A native hormone is usually present only in trace amounts in the feces or can be totally absent [32–34]. Thus, using a native hormone assay for non-invasive samples, for example, feces, urine, is not appropriate and may lead towards dubious results. This explains the higher levels of cortisol metabolite in our study when compared to Vaz et al. (2017) [21]. Further, Vaz et al. (2017) [21] measured corticosterone whereas cortisol is the predominant glucocorticoid released in mammals [35–37].

In the current study, changes in adrenocortical activity in captive Indian leopards was detected by measuring concentration of a cortisol metabolite (5α-pregnane-3β,11β,21-triol-20-one), and the same assay has been used for African leopards [9]. Thus, we directly compared fGCM levels in captive individuals between the African (*pardus* subspecies) and the Indian subspecies (*fusca* subspecies), and found the mean fGCM value to be 36 times higher in Indian leopards than their African counterparts [9]. Such a huge difference in fGCM values between the Indian and the African subspecies can be due to multiple factors, ranging from differences in local environment to physiological differences at a subspecies level. For example, a study on white crowned sparrows (*Zonotrichia leucophrys pugetnesis, Zonotrichia leucophrys gambelli and Zonotrichia leucophrys nuttalli*) demonstrated that individuals breeding at higher altitude had increased stress hormone levels when compared to the subspecies which bred at lower altitude [38]. In contrast, Gonzalez at el. (1981)[39] demonstrated subspecies-level differences in plasma cortisol concentrations for squirrel monkeys (*Saimiri sciureus sciureus* and *Saimiri sciureus albigena*) when reared under identical laboratory conditions. In the current study, such differences in fGCM levels were not observed between free-ranging Indian and African subspecies. The fGCM values of the free-ranging Indian leopards lie within the range of cortisol values reported for the free-ranging African subspecies [9]. However, the sample size for free-ranging Indian leopards was significantly lower in our study. The scat samples collected from free-ranging leopards were for the purpose of validating the fGCM assay through generating parallelism curves, and any further insights on the fGCM patterns of free-ranging leopards will warrant future investigations.

Interestingly, there were no significant differences in fGCM levels between captive male and female leopards for both the African [9] and the Indian subspecies (the current study). Sex-specific differences in cortisol are not prevalent in other carnivores [40, 41], as well. A study on tigers [41] showed no sex-specific differences in fGCM levels, however, pregnant females had significantly higher fGCM levels than males and non-pregnant females. Thus, the effect of sex type on fGCM levels can be very variable, and can be further influenced by several physical and social factors, for example, social rank, age, life history, and reproductive status [42]. In the current study, the sampled leopards were all adults with no breeding history and were held in social isolation (one leopard per cage). Such lack of differences in physical and social make-up may possibly explain the similar fGCM profiles for both the sex types within the sampling population.

The results from our study indicate a statistically significant effect of enrichment on the fGCM values of captive leopards. In the current study, the leopards in category ‘B’ had the highest mean (season-wise and pooled across seasons, as well) fGCM values when compared to the other two categories. In contrast, leopards in category ‘C’ had the lowest mean (season-wise and pooled across seasons) value of stress hormone metabolites. Categories ‘B’ and ‘C’ were similar in quality having only elements of active enrichment (earthen floor, raised platforms). The only difference was that category ‘B’ had the smallest cage size and was provided with air coolers during summer. In contrast, category ‘C’ had medium size cages (48.94% larger than ‘B’) with no air coolers during summers and category ‘A’ had the largest cage size (78.07% larger than ‘B’) with several passive enrichment elements, as well. Thus, high fGCM values of leopards in category ‘B’ indicated smaller cage size to be an important active, enrichment element that may contribute towards increased levels of stress hormone metabolite levels in the leopards under the ‘B’ regime. Similar results have been documented in other species, as well, where one of the elements had a stronger physiological effect than the rest within a given enrichment regime. Lapinski at el. (2019) [43] showed that cage size had no effect on glucocorticoid metabolite levels in fox (*Vulpes vulpes*). However, gnawing stick, another type of active enrichment, lowered the salivary glucocorticoid metabolite levels of captive fox, from 4.65±0.98 to 3.70±1.01 ng/ml. In the current study, other variables, for example, keeper’s attitude towards the leopards, occurrence of diseases in captivity, frequency of visitors, have not been measured across different enrichment categories, which could possibly impact the fGCM levels and needs further systematic analyses.

Studies have documented that physiological effect of enrichment elements is quite diverse and usually varies from species to species, and even across different individuals of the same species. For example, a study on the Asian elephants [44] showed differences in fGCM levels across individuals, though, all were provided with similar kinds of structural enrichment. This pattern is also evident in our study, where within category ‘A’ and category ‘B’, all individuals had high variations in fGCM levels. This is more evident in category ‘B’, where individual variations are more prominent irrespective of the differences across seasons. Interestingly, category ‘B’ also had very high values of mean fGCM than the other two categories. Literature has documented that high levels of stress hormone often generates a lot of individual variations. For example, a study in barn swallows [45] showed that under high predation rate, the GC levels increased, and this was also accompanied with a high degree of individual variations in the hormone levels. Thus, to understand the physiological wellbeing of a target species, systematic monitoring at an individual level needs to be conducted.

The current study is the first of its kind to include passive-enrichment elements in understanding the wellbeing of the captive leopards. Category ‘A’ was the only regime where passive-enrichment elements, for example, natural sounds, sound proof glasses to filter visitors’ noise and artificially maintained lower ambient temperature, were provided along with active-enrichment components, earthen floor and relatively large cage size. However, there were no significant differences in fGCM values between categories ‘A’ and ‘C’ during the summer, where ‘C’ is the regime with active-enrichment elements only. Thus, at a physiological level, the combination of active and passive elements in category ‘A’ had a similar effect on fGCM levels when compared to active enrichment regime in category ‘C’. To the best of our knowledge, no study has shown the combined effect of active and passive enrichment elements on the stress physiology of captive leopards.

The current study did not find any significant effect of seasons on fGCM levels, except for the leopards in category ‘A’. fGCM levels of leopards in category ‘A’ were significantly different between the summer and winter seasons, with higher levels during winter. However, we speculate that the higher fGCM levels in winter is not a ‘season effect’ but can be attributed to the presence of two old (>19 years) individuals in category ‘A’. Both the individuals showed high values during winter and eventually died after a span of 8-10 months. The effect of age on physiological stress levels is dynamic, and is very context-specific. For example, several studies found that old age is related to higher fGCM levels but similar studies in the same species reported a contrasting result, demonstrating lower levels of fGCM in old age animals [46–48]. Thus, in our study, we speculate that the higher cortisol levels in the two individuals in category ‘A’ can be because of old age but such speculation needs to be corroborated with long-term, longitudinal fGCM profiles of captive leopards across different life-history stages.

In conclusion, the current study demonstrated that enrichment, in particular the size of the cage, influenced the fGCM levels of the Indian leopards within our sampling population. However, our study also demonstrated that physiological response is quite diverse, showing huge variations in fGCM levels across individuals, and even for the same individual between the summer and winter seasons. Thus, zoo management programmes should focus on including a large number of covariates, including both individual-(for example, sex type, age, life-history stage, time since captivity, social rank, health and disease status of the animal) and environment-specific (for example, types of enrichment element, diet regime, presence of conspecific individuals, seasons, visitors’ frequency, and keeper’s attitude) factors to understand the physiological wellbeing of the target species within a captive environment. Further, the current study used an fGCM assay method that has previously been validated for African leopards and standardized the same for the Indian leopards, as well, under both captive and free-ranging conditions. Such applications of the same assay method for different sub-species in geographically different regions allows for direct comparisons of endocrine profiles within or across a taxonomic group. In terms of management practices, the current study provided a standardized fGCM assay to monitor the wellbeing of Indian leopards and demonstrated the overall importance of cage size as an enrichment element for the sampled leopard population. The findings of the current study will contribute towards developing informed management policies for conservation of the Indian leopards.

## Acknowledgements

The authors would like to thank the zoo directors Shri. RK Sahu and Dr. Pratyush Patankar for giving the permission to collect scat samples from Kankaria and Baroda zoos, respectively. The authors also thank Shri. Shyamal Tikadar (Principal Chief Conservator of Forests, Forest Department, Gujarat, India) for providing permit to collect scats in protected areas, and Krunal Trivedi, Surat Nature Club for cooperation and support during sample collection from wild leopards. The project was supported through Ahmedabad University’s start-up grant awarded to RG. NP and CD acknowledges the Integrated master’s programme of Ahmedabad University for providing an opportunity to conduct the research as a part of their master’s dissertation project.

---

Supplemental Information (for review purpose only)

**Table 1:** Sampling details of leopards in Kankaria and Sayajbaug zoos

| Enrichment regime | Number of samples per individual | Total number of samples | Season |
| --- | --- | --- | --- |
| A | 3 | 12 | Summer |
| A | 3-7 | 21 | Winter |
| B | 3-4 | 14 | Summer |
| B | 2-5 | 13 | Winter |
| C | 5-7 | 36 | Summer |
| C | 4-6 | 31 | Winter |

**Table 2:** Comparative overview of fecal glucocorticoid metabolite (fGCM) levels in Indian leopards from the current study and in African leopards from published data, and fecal glucocorticoid (fGC) levels in Indian leopards from published literature.

| fGCM levels (Mean±SE, range) (µg/g dry wt of feces) in Indian leopards (current study) |  | fGC levels (Mean±SE) (µg/g dry wt of feces) in Indian leopards (Vaz *et al.* 2017) |  | fGCM levels (Mean value range) (µg/g dry wt of feces) in African leopards (Webster *et al.* 2018) |  | Remarks |
| --- | --- | --- | --- | --- | --- | --- |
| Captivity | Wild | Captivity | Wild | Captivity | Wild | Webster *et al.* and the current study used the same hormone assay for |
| 10.40±3.87, | 0.96± 0.16, 0.2-1.93 | 0.27±0.013 | NA | 0.29-0.39 | 0.02-6.09 |  |
| 0.2-108.63 |  |  |  |  |  | measuring<br>fGCM levels |

## Notes

### Competing Interest Statement

The authors have declared no competing interest.

